# Cytomegalovirus infection is a risk factor for TB disease in Infants

**DOI:** 10.1101/222646

**Authors:** Julius Muller, Rachel Tanner, Magali Matsumiya, Margaret A. Snowden, Bernard Landry, Iman Satti, Stephanie A. Harris, Rachel Tanner, Matthew K. O’Shea, Lisa Stockdale, Leanne Marsay, Agnieszka Chomka, Rachel Harrington-Kandt, Zita-Rose Manjaly Thomas, Vivek Naranbhai, Elena Stylianou, Stanley Kimbung Mbandi, Mark Hatherill, Gregory Hussey, Hassan Mahomed, Michele Tameris, J. Bruce McClain, Thomas G. Evans, Willem A. Hanekom, Thomas J. Scriba, Helen McShane, Helen A. Fletcher

## Abstract

Immune activation is associated with increased risk of tuberculosis (TB) disease in infants. We performed a case-control analysis to identify drivers of immune activation and disease risk. Among 49 infants who developed TB disease over the first two years of life, and 129 matched controls who remained healthy, we found the cytomegalovirus (CMV) stimulated IFN-γ response at age 4-6 months to be associated with CD8+ T-cell activation (Spearman’s rho, p=6×10^−8^). A CMV specific IFN-γ response was also associated with increased risk of developing TB disease (Conditional Logistic Regression, p=0.043, OR 2.2, 95% CI 1.02-4.83), and shorter time to TB diagnosis (Log Rank Mantel-Cox p=0.037). CMV positive infants who developed TB disease had lower expression of natural killer cell associated gene signatures and a lower frequency of CD3-CD4-CD8-lymphocytes. We identified transcriptional signatures predictive of risk of TB disease among CMV ELISpot positive (AUROC 0.98, accuracy 92.57%) and negative (AUROC 0.9, accuracy 79.3%) infants; the CMV negative signature validated in an independent infant study (AUROC 0.71, accuracy 63.9%). Understanding and controlling the microbial drivers of T cell activation, such as CMV, could guide new strategies for prevention of TB disease in infants.

## INTRODUCTION

There are an estimated 1 million cases of childhood tuberculosis (TB) each year and in 2015, 210,000 children died of TB ^5^. Children with TB are difficult to diagnose and treat and are at risk of severe disease ^6^. The need for improved strategies to control childhood TB has led to studies to identify risk factors for TB disease in children. The longitudinal data collected during infant TB vaccine efficacy trials in South Africa with the vaccines BCG and Modified Vaccinia Virus Ankara expressing Antigen 85A (MVA85A) ^1, 7^ have enabled the identification of correlates of TB disease risk in infants ^3, 8^. Using samples collected at enrolment from infants into the MVA85A efficacy trial in the Western Cape Province of South Africa ^1^, we reported that CD4+ T-cell activation in 4-6 month old infants and adolescents, measured as HLA-DR expression, was associated with increased risk of TB disease over the following 3 years of life ^8^. The consequences of chronic T-cell activation are well-described in HIV and include risk of acquisition of infection, risk of progression from infection to disease, increased risk of non-communicable disease and aging ^10-14^. Recognition that T-cell activation is a feature of HIV immunopathogenesis has guided the development of new interventions for management of HIV ^15^. Improved understanding of the causes and impact of T-cell activation on TB disease risk, could transform future approaches to protect against TB. Such approaches could include the use of vaccines, antibiotics or antivirals to reduce the burden of chronic microbial drivers of T cell activation.

Sustained T-cell activation and dysfunction of antigen specific T-cells can result from chronic exposure to antigen from persistent viral or bacterial infections ^9^. Human cytomegalovirus (CMV) and Epstein-barr 1 virus (EBV) are known drivers of T-cell activation ^11,14^ and with more than 95% seroprevalence in adults, South Africa is amongst the highest CMV burden countries in the world (reviewed by Adland et. al., ^16^). Recent epidemiological evidence supports a role for CMV in the aetiology of TB ^17-21^. Here, we report results from a case–control analysis of infants from the MVA85A efficacy trial ^1^ which aimed to understand the drivers of immune activation associated with TB risk in infants.

## RESULTS

### Activated CD8+ T-cells are associated with TB disease risk and correlate with cytomegalovirus (CMV)-specific IFN-γ response

BCG-vaccinated case and control infants who were enrolled in an efficacy trial of the candidate TB vaccine MVA85A were included in this study ^1^ (Figure 1). HIV and *M.tb* uninfected infants without known TB exposure were randomized at 16-24 weeks of age to receive a single intradermal dose of MVA85A or placebo (Candin™, a candida skin test antigen)^1^. In contrast to our previous analysis which focused on the sample collected at enrolment (Day 0), samples collected from two time points, Day 0 (D0) and Day 28 (D28) following MVA85A or placebo were combined for testing of both cellular parameters and transcriptional signatures associated with risk of TB disease. We have shown that the correlates of risk measured in our previous study were not affected by intervention (MVA85A or placebo) ^8^. Only infants for whom a sample was available at both D0 and D28 were included in the analysis (Figure 1).

**Figure 1.**
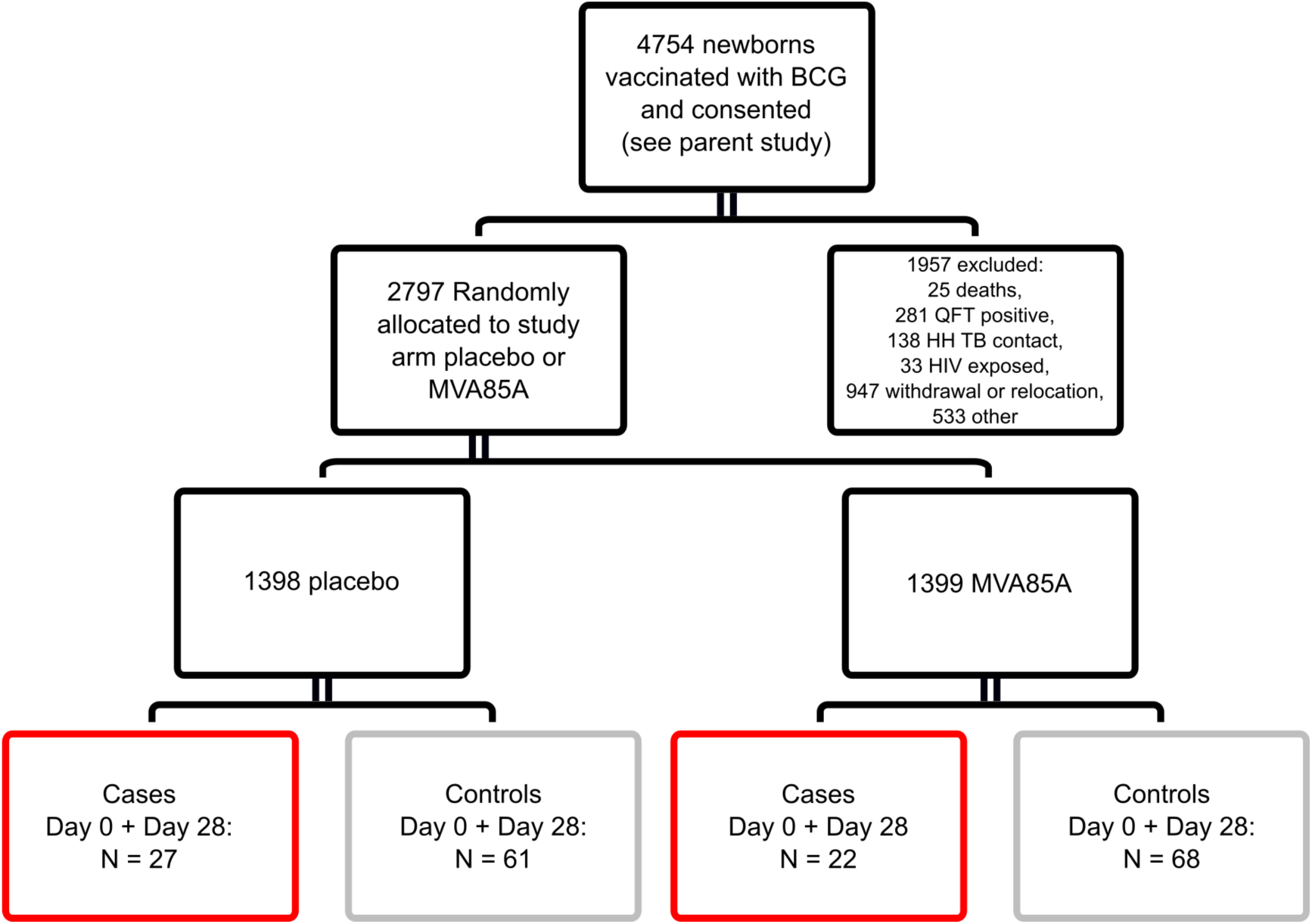
Study design for immune correlates analysis. Infants who were enrolled in an efficacy trial of the candidate TB vaccine MVa85A were included in this study ^1^. Infants were randomized at 16-24 weeks of age to receive a single intradermal dose of MVA85A or placebo (Candin™, a candida skin test antigen)^1^. Boxes indicate the number of case infant (red) or control infant (grey) samples available for combined Day 0 (D0) and Day 28 (D28) analysis. Analysis was restricted to infants where a frozen PBMC sample was available, live cells in PBMC were >50% (or PHA IFN-γ ELISPOT >=1000 SFC/million) and to infants where a sample was available for analysis from both the D0 and D28 time points. Control infants were excluded if the corresponding matched case was not in the analysis.

Previously, we identified an association between TB disease risk over the first two years of life and the frequency of activated HLA-DR+ CD4+ T-cells at age 4-6 months ^8^. We also found that the magnitude of BCG-specific IFN-γ expressing cells and levels of anti-Ag85A IgG were associated with reduced risk of TB disease ^8^. In our present analysis, with combined Day 0 and Day 28 samples from 49 cases and 129 controls (Figure 1), we confirmed our previous finding of TB risk associating with HLA-DR+ CD4+, BCG-specific IFN-γ and anti-Ag85A IgG (Table 1). In addition, we found frequencies of HLA-DR+ CD8+ T-cells to be associated with TB risk (conditional logistic regression (CLR) OR 1.34, 95% CI 1.08-1.67, p = 0.008, FDR 0.092) (Table 1). The magnitude of PPD-stimulated IFN-γ-expressing cells measured by ELISpot were also associated with reduced TB risk (OR 0.71, 95% CI 0.51-0.98, p = 0.037, FDR 0.2) (Table 1).

**Table 1.** Conditional Logistic Regression of combined Day 0 plus Day 28 infant samples

| Estimated Odds-Ratio (OR) of TB disease from a Conditional Logistic Regression of D0 + D28 immunological variable |  |  |  |  |  |  |  |
| --- | --- | --- | --- | --- | --- | --- | --- |
| Quantitative variable | N | Case | Est OR* | 95% CI | P value | FDR value | AUROC |
| CD3+ T cell | 329 | 88 | 0.98 | 0.75-1.28 | 0.86 | 0.94 |  |
| CD4+ T cell | 329 | 88 | 0.79 | 0.6-1.03 | 0.087 | 0.29 |  |
| HLA-DR+ CD4 T cell | 329 | 88 | 1.59 | 1.3-2 | <0.0001 | 0.002 | 0.64 |
| CD4+CD25+CD127- | 329 | 88 | 0.84 | 0.62-1.12 | 0.23 | 0.59 |  |
| CD8+ T cell | 329 | 88 | 1.23 | 0.98-1.53 | 0.074 | 0.28 |  |
| HLA-DR+ CD8+ T cell | 329 | 88 | 1.34 | 1.08-1.67 | 0.008 | 0.092 | 0.59 |
| CD14+CD16+ | 329 | 88 | 0.93 | 0.73-1.18 | 0.57 | 0.82 |  |
| CD14+CD16- | 329 | 88 | 0.99 | 0.78-1.27 | 0.96 | 0.96 |  |
| CD19+ B cell | 328 | 88 | 1.13 | 0.87-1.4 | 0.39 | 0.69 |  |
| BCG MGIA | 146 | 50 | 0.82 | 0.52-1.29 | 0.39 | 0.69 |  |
| 85A ELISpot | 324 | 91 | 0.87 | 0.63-1.2 | 0.39 | 0.69 |  |
| BCG ELISpot | 333 | 94 | 0.75 | 0.57-0.98 | 0.036 | 0.2 | 0.56 |
| PPD ELISpot | 270 | 79 | 0.71 | 0.51-0.98 | 0.037 | 0.2 | 0.57 |
| TB10.3/10.4 ELISpot | 270 | 79 | 0.96 | 0.74-1.23 | 0.72 | 0.83 |  |
| EBV ELISpot | 290 | 84 | 1.05 | 0.82-1.35 | 0.7 | 0.83 |  |
| CMV ELISpot | 319 | 91 | 1.01 | 0.8-1.23 | 0.9 | 0.94 |  |
| FLU ELISpot | 289 | 84 | 1.08 | 0.85-1.37 | 0.55 | 0.82 |  |
| GAM.DEL (putative) | 329 | 88 | 1.16 | 0.91-1.5 | 0.23 | 0.59 |  |
| NK 16neg (putative) | 329 | 88 | 0.96 | 0.63-1.4 | 0.63 | 0.82 |  |
| NK 16pos (putative) | 329 | 88 | 0.86 | 0.66-1.12 | 0.26 | 0.6 |  |
| CD14+CD16+ /CD3+ | 329 | 88 | 0.92 | 0.72-1.18 | 0.52 | 0.82 |  |
| CD14+CD16- /CD3+ | 329 | 88 | 0.96 | 0.74-1.22 | 0.68 | 0.83 |  |
| Anti-Ag85A IgG | 297 | 87 | 0.79 | 0.63-0.99 | 0.043 | 0.2 | 0.59 |

We analyzed CMV and EBV-specific IFN-γ ELISpot 1 responses for evidence of an association between viral infection and T-cell activation in our infant cohort. Frequencies of activated CD8+ T-cells correlated with the magnitude of the CMV-specific IFN-γ ELISpot response (Spearman’s rho p = 6 × 10^−8^, Figure 2A), suggesting that CMV infection is associated with CD8+ T-cell activation in this infant cohort. A network representation of positively correlating cell populations ^4^ (spearman rho p-value <0.05) revealed 3 clusters dominated by activated CD8+ and activated CD4+ T-cells with CMV-specific IFN-γ ELISpot response, CD3+ T-cells with Epstein–Barr virus (EBV)-specific IFN-γ ELISpot response and monocytes with B-cells (Figure 2B).

**Figure 2.**
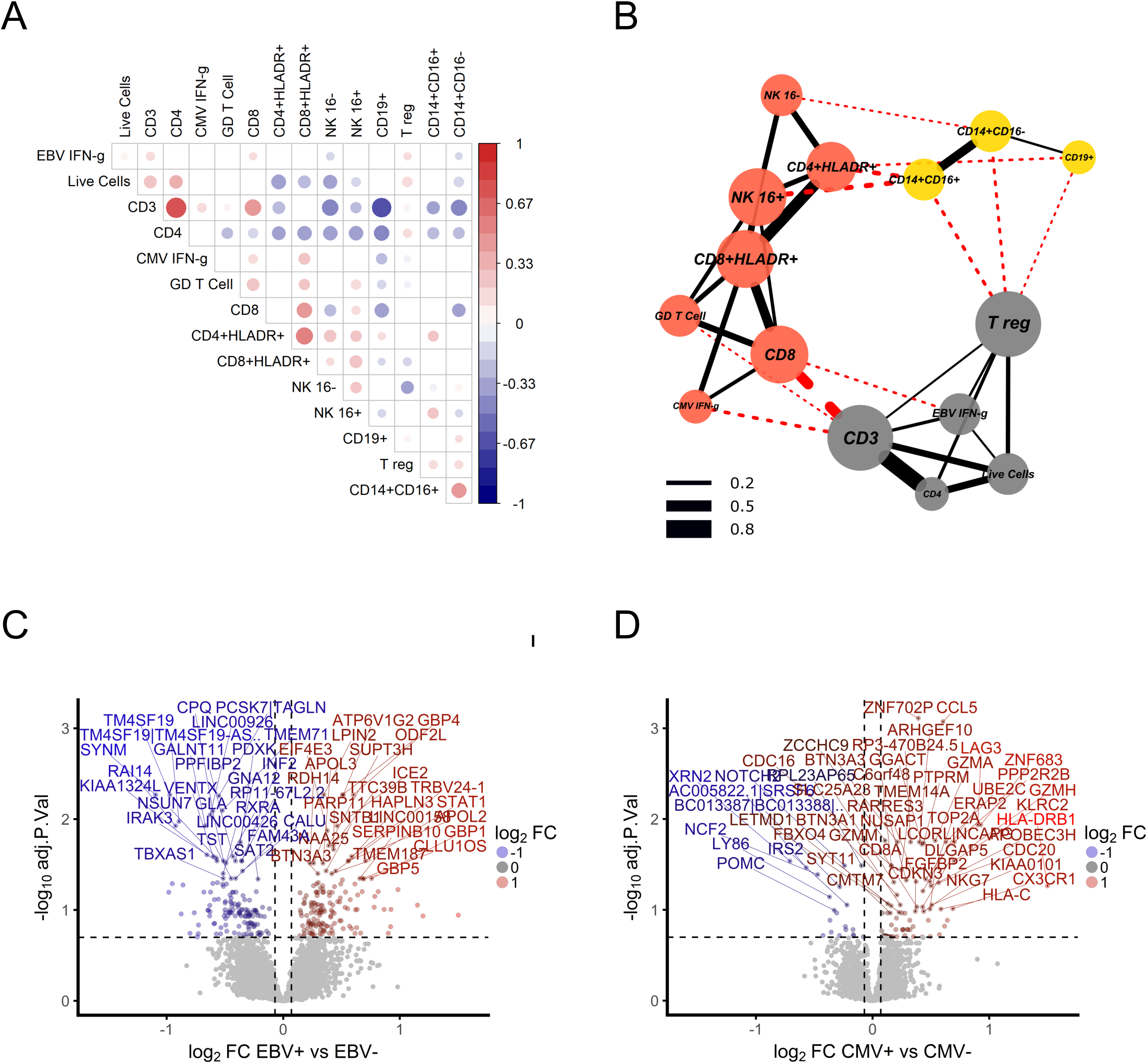
CMV is associated with CD8 T cell activation in South African infants. A) Correlation matrix of significantly (spearman rho p <0.05) correlated cell populations and IFN-γ ELISpot responses to EBV and CMV. The magnitude of the CMV specific IFN-γ ELISpot response correlated with the frequency of activated CD8+ T cells. B) Network of positively correlating cell populations (spearman rho p <0.05) showing 3 clusters dominated by activated T cells with CMV, CD3+ T cells with EBV and monocytes with B cells (node colour indicates cluster membership^4^). Red lines indicate between cluster correlations and black lines within cluster correlations. Line width indicates the correlation coefficient. C) Volcano plot showing magnitude and significance of differential expression between EBV+ and EBV- infants and D) CMV strongly positive (ELISpot >100 SFC/million) and CMV negative infants. The top 50 significant genes are labelled and horizontal and vertical dashed lines indicate 20% FDR and 5% change in gene expression respectively.

EBV had a strong effect on the blood transcriptome, with 296 genes differentially expressed between infants with positive and negative EBV responses (Figure 2C). CCL8, CXCL10 and IFIT3 had the greatest fold increase in expression in EBV+ infants indicating strong induction of a Type I/II IFN response, although we did not see a correlation between EBV ELISpot response and T cell activation in this study (Supp Table 1).

The impact of CMV on the blood transcriptome was smaller, with only 14 genes differentially expressed between CMV positive and negative infants (ELISpot >17 SFC/million), although there were 103 differentially expressed transcripts between CMV strongly positive (ELISpot >100 SFC/million) and negative infants (Figure 2D and Supp Tables 2A and B). Differentially expressed transcripts included HLA-DRB1, ZNF683, LAG3 and KLRC2 (NKG2C). ZNF683 is an important paralog of PRDM1 and together with CCL5 has been shown to be highly induced in human CMV- specific CD8+ T-cells when compared to naïve CD8+ T-cells ^22^. LAG3 is expressed on activated CD8+ and CD4+ T-cells and LAG3+CD8+ T-cells are found in CMV infection ^22,23^. NKG2C+ natural killer (NK) cells are expanded in response to CMV infection although CMV infection can also induce the expression of NKG2C on CD8+ T-cells ^24-26^.

Expression levels of transcripts from total peripheral blood mononuclear cells (PBMC) of infants with a CMV ELISpot response (>100 SFC/million) were correlated with expression levels of transcripts published from CD8+ T-cells isolated from adults at different stages of CMV infection ^22^. The highly significant correlation of infant transcripts with CMV virus specific CD8+ T-cell transcripts from adults confirms the prominence of an activated CD8+ T-cell response in unstimulated PBMC of CMV ELISpot positive infants (Additional Figure 1 A and B).

Interestingly, there was almost no overlap in differentially expressed genes between CMV and EBV positive infants with only seven genes significantly differentially expressed in both comparisons at an FDR of 20% (Additional Figure 2).

Our cellular data suggests that CMV IFN-γ responses are associated with activated HLA-DR+ CD8+ T-cells and transcriptional analysis supports the presence of an activated CD8+ T-cell phenotype amongst total, unstimulated PBMC from CMV IFN-γ ELISpot positive infants.

### CMV antigen-specific T-cell responses, measured up to 3 years prior to detection of TB, associate with risk of TB disease in infants

CMV is associated with increased risk of HIV infection and disease progression ^11,12,14^. To determine if viral infection is associated with TB risk in our infant cohort, we analyzed CMV- specific and EBV-specific IFN-γ ELISpot responses in cases and controls. Infants dual positive for both EBV and CMV (n=3) were excluded from the analysis. A CMV specific IFN-γ ELISpot response (response >17 SFC/million at either time point), measured up to 3 years prior to TB detection, was associated with increased risk of TB disease (CLR, p=0.043, OR 2.22 95% CI 1.02-4.83) (Figure 3A). EBV alone was not associated with risk (Figure 3B), although a combined CMV and EBV response was associated with increased risk of TB disease (CLR, p=0.025, OR 2.3 95% CI 1.11-4.79) (Figure 3C). To further explore this association, we analyzed time to TB diagnosis in TB cases. Infants with a positive CMV or CMV/EBV+ ELISpot response developed TB earlier than negative infants (Log Rank Mantel-Cox p=0.037, Figure 3D and 3E).

**Figure 3.**
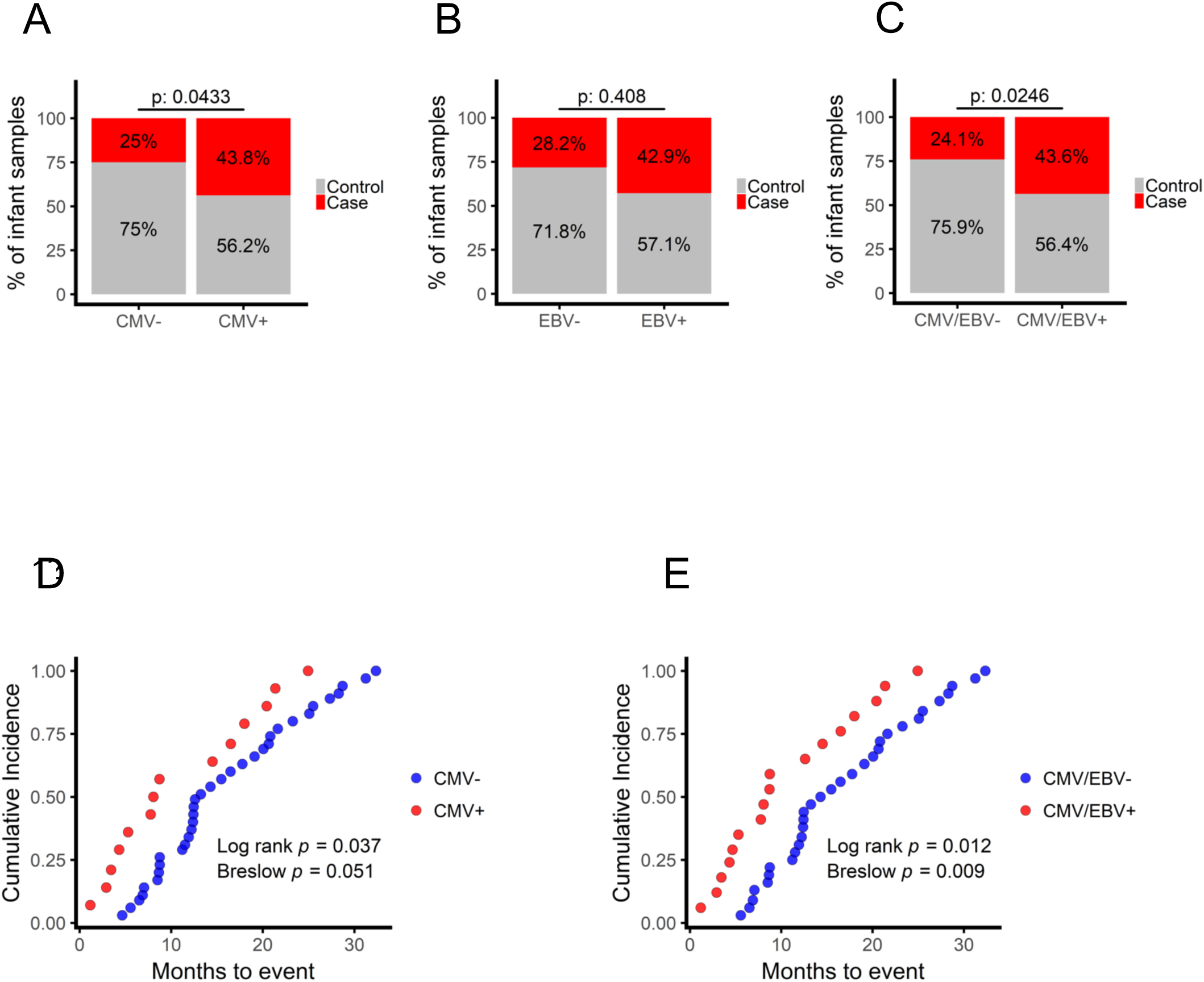
CMV+ infants are at increased risk of developing TB disease. A) We saw a higher proportion of case (red) infants among CMV+ (n = 18/32) when compared to CMV- infants (n = 35/140). B) There was no significant enrichment for cases among EBV positive infants (n = 3/7 compared to n = 46/162) although Infants positive for either CMV or EBV (C) were at increased risk D) CMV+ infants (red) develop TB disease earlier in follow-up when compared to CMV- infants (blue) and E) Infants positive for either CMV or EBV develop TB disease earlier than CMV/EbV- infants.

We analyzed data by vaccine group and saw no evidence that CMV positive or CMV/EBV positive infants immunized with MVA85A were at greater or lower risk of TB disease when compared to CMV negative infants immunized with MVA85A (p = 0.8355 for CMV and p = 0.9177 for CMV/EBV).

A CMV specific IFN-γ response measured at 4-6 months of age, up to 3 years before disease is detected, was a risk factor for the development of TB disease in South African infants and this risk was greatest during the first 10 months of follow-up.

### Transcriptional evidence of activated T-cells, Type I IFN responses and NK cells in infants up to 3 years prior to detection of TB

Because CMV ELISpot positive infants were at greater risk of disease, we stratified transcriptome data by CMV status to identify transcripts able to classify case and control infants. When CMV+ and CMV- infants were analyzed together, 16 genes were significantly differentially expressed between cases and controls (Figure 4A and Supp Table 3). LAG3 and VCAM1, known markers of T-cell activation ^23,27,28^, had the greatest fold increase in expression in case infants. To test our ability to classify infants into cases or controls we split samples randomly into a 70% training set and 30% test set and trained an artificial neural network (Additional Figure 3A). The process was repeated fifty times with random splits into training and test set (bootstrapping) and AUROC and accuracy were recorded to assess predictive stability (Additional Figure 4). When all infants were included in the analysis we could classify infants into cases and controls with an average accuracy of 67% (AUROC 0.77, 95% CI 0.69-0.85, Additional Figure 6 A). Prediction accuracies were comparable when alternative, widely used classification algorithms were used (Additional Figure 3B).

**Figure 4.**
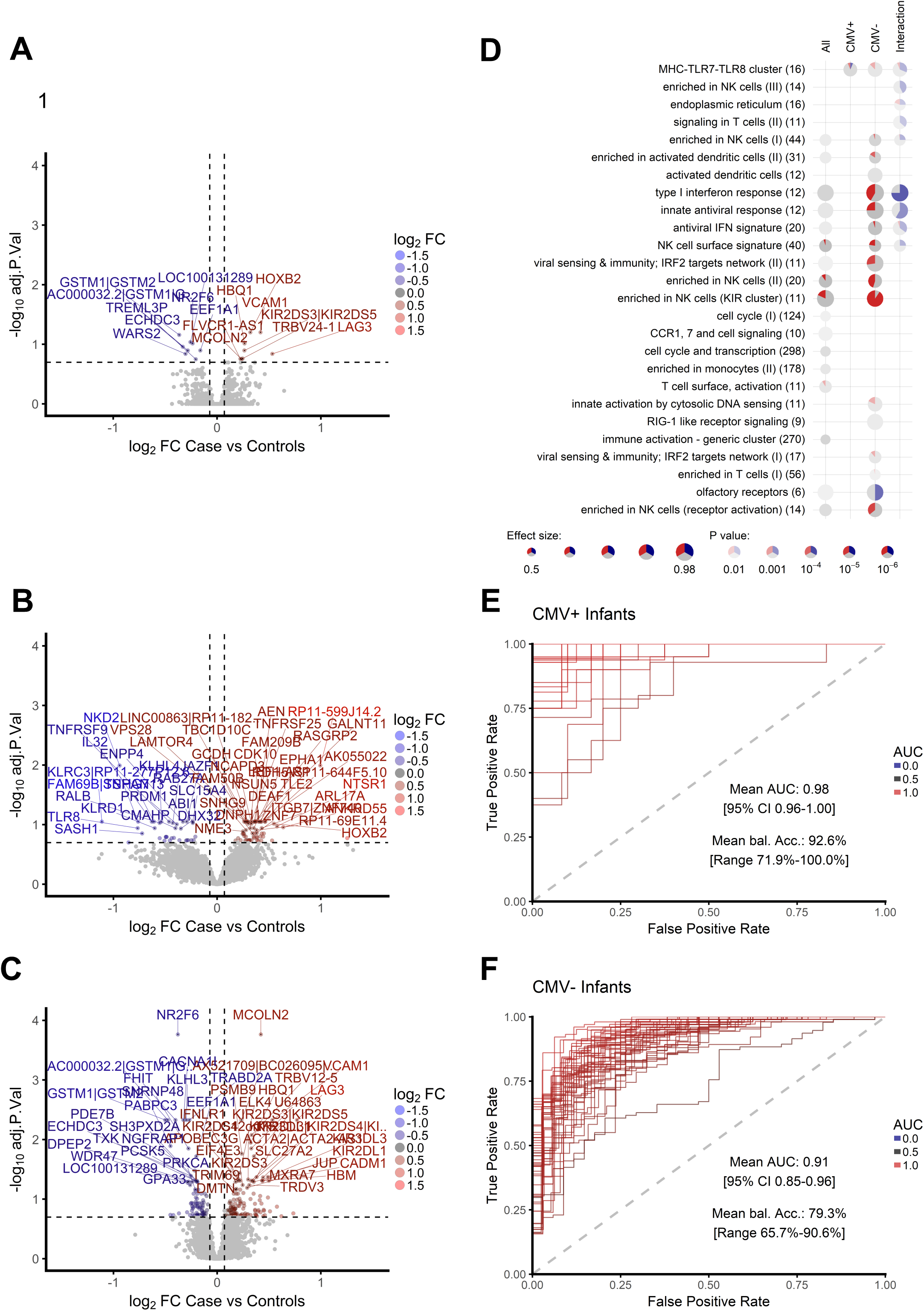
Transcriptomic correlates of risk of TB disease are different in CMV+ and CMV- infants. Volcano plots showing magnitude and significance of differential expression between all case and control infants A), CMV+ case and control infants B) and C) CMV- case and control infants. The top 50 significant genes are labelled and horizontal and vertical dashed lines indicate 20% FDR and 5% change in gene expression respectively. D) Enriched modules for differential expression in case and control infants amongst all, CMV+ and CMV- infants. Each row contains one module with the number of genes indicated. Each significantly enriched module at a p-value < 0.05 is shown as a pie chart. The size of the pie corresponds to the AUROC in the cerno test, and intensity of the colour corresponds to the enrichment q-value. The red and blue colour indicates the amount of significant up and down regulated genes respectively, grey color indicates the remaining not significant genes within the category. The interaction term evaluates the statistical difference between changes in CMV+ and CMV- infants. E) AUROC of the classification performance of the artificial neural network model which was trained using approximately 70% of the data and risk of TB was predicted on the withheld 30% of the data (Additional Figure 3 A). The process was repeated fifty times with random splits into training and test set (bootstrapping) and the AUROC was recorded for each round for E) CMV+ and F) CMV- infants respectively.

In CMV+ healthy infants who developed TB in the following 3 years (cases), the NK cell associated cytokine IL32 and the NK cell-specific lectin-like receptors KLRC1 and KLRC3 were among transcripts with the greatest decrease in fold-change of expression and the highest predictive power (Figure 4B, Additional Figure 4 and Supp Table 4). IL32 enhances maturation of monocytes to macrophages and has been shown to be important for protection against *Mycobacterium tuberculosis* (*M.tb*) ^29^ In our cellular analysis we observed decreased frequencies of CD3-CD8-CD4- (triple negative) CD16- and CD16+ natural killer cells in infants who develop TB disease in the next 3 years when compared to CMV+ infants who do not develop disease (Additional Figure 6B). In CMV- case infants, elevated expression of T-cell activation markers, including LAG3 and VCAM1, markers of a type I/II IFN response including IFIT3, and enhanced expression of a broad range of both activated and inhibitory KIR receptors including KIR2DL1, KIR2DL3, KIR2DL4, KIR2DL5A, KIR2DS3, KIR2DS5, KIR3DL1 and KIR3DL3 were observed (Figure 4C and Supp Table 5). Modular pathway analysis showed enrichment for NK and KIR cluster genes and a type I/II IFN antiviral immune response in CMV- case infants when compared to controls (Figure 4D). However, we saw no evidence of increased NK cell frequency in infants who develop TB disease in our cellular analysis (Additional Figure 6B).

TB risk associated transcripts and immune pathways were different among CMV+ and CMV- infants, as highlighted by a modular pathway analysis which uses an interaction term to compare pathways associated with TB risk among CMV+ and CMV- infants (Figure 4D and Supp Table 6).

When infants were stratified by CMV status we were able to classify cases and controls within the CMV+ cohort with 92% average balanced accuracy and within the CMV- cohort with 89% average balanced accuracy with an average AUROC of 0.98 (95% CI 0.96-1) and 0.9 (95% CI 0.85-0.96), respectively (Figure 4 E and F and Additional Figures 4).

To validate the classifier signatures, we used raw data from an independent cohort of 10-week-old South African infants vaccinated with BCG at birth and unknown CMV status (GEO gene set GSE20716, Fletcher *et. al.* ^3^) Infants were enrolled into an efficacy trial of intradermal or percutaneous delivery of Japanese BCG at birth ^7^. A nested correlates of risk study was performed using blood from 10-week-old healthy infants who developed TB disease within the next two years ^3, 30^. We sought to remove technical differences between the two array data sets to enable validation of classifier signatures (Additional Figure 7A), however, we could not fully normalize expression between the two data sets, most likely due to the age difference between infants in the two cohorts (2-3 months in GSE20716 and 4-6 months in the MVA85A efficacy trial) (Additional Figure 7B). Despite these cohort differences and the loss of more than half of the genes in our signature due to microarray platform differences, we were able to predict risk of TB using our CMV- classifier with an accuracy of 63.9% and an AUROC of 0.71 (95% CI 0.63-0.79, Additional Figure 7C). Next, we attempted to improve accuracy by prediction and removal of infants with suspected CMV infection. The number of CMV positive infants with more than one predicted positive sample was low (n = 5, Additional Figure 7C) and excluding these infants lead to a slight but not significant improvement in balanced accuracy of 65.6% and an AUROC of 0.74 (Additional Figure 7E).

Finally, we looked for enrichment of transcripts from the 16- peripheral blood gene correlate of risk (CoR) signature associated with progression to TB disease in *M.tb* infected adolescents^2^ among our CMV+ and CMV- infants. This adolescent CoR signature was significantly enriched among case infants when compared to controls and the enrichment was strongest amongst CMV- infants (Additional Figure 8A). However, the adolescent CoR signature was not able to accurately classify infants into cases and controls (Additional Figure 8B). Including the CoR score in our infant network analysis we show that the CoR score correlates with the frequency of inflammatory monocytes and with our CMV- classifier signature (Additional Figure 8C). Our CMV+ classifier signature is positively correlated with T cell frequency and negatively correlated with monocyte and CD16+ NK cell frequency.

A network representation of positively correlating cell populations (Spearman’s rho p-value <0.05) revealed 3 major clusters with the adolescent CoR and infant CMV- classifier signature clustering together with inflammatory and classical monocytes (Additional Figure 8D).

These findings further support distinct immunological correlates of risk of TB disease in CMV+ and CMV- infants. Taken together our data show increased T-cell activation, KIR receptor signaling and type I IFN response in CMV- case infants and decreased NK cell associated transcripts in CMV+ case infants when compared to their respective controls. Furthermore, we identified transcriptomic signatures, which can identify infants with risk of TB disease with high accuracy in CMV+ and CMV- infants and were able to validate the CMV- biomarker signature in an independent study with moderate accuracy.

### T-cell activation associates with lower mycobacterial antigen specific immune response following immunization with MVA85A and BCG

To assess the impact of T-cell activation and CMV on the ability to mount an antigen specific response following immunization with MVA85A, we examined mycobacterial antigen specific IFN-γ responses and anti-Ag85A IgG in MVA85A immunized infants at D28 following immunization with MVA85A. IFN-γ responses to Ag85A, measured on D28, were inversely correlated with activated CD4+ and CD8+ T-cell frequencies (Figure 5A and B). There was a trend towards lower mycobacterial antigen specific IFN-γ responses and lower anti-Ag85A IgG in CMV+ when compared to CMV- infants (p = 0.058, Mann-Whitney U test) and Anti-Ag85A IgG was inversely correlated with CMV ELISpot response in MVA85A immunized infants (Spearman’s Rho, p = 0.026, Figure 5 C and D).

**Figure 5.**
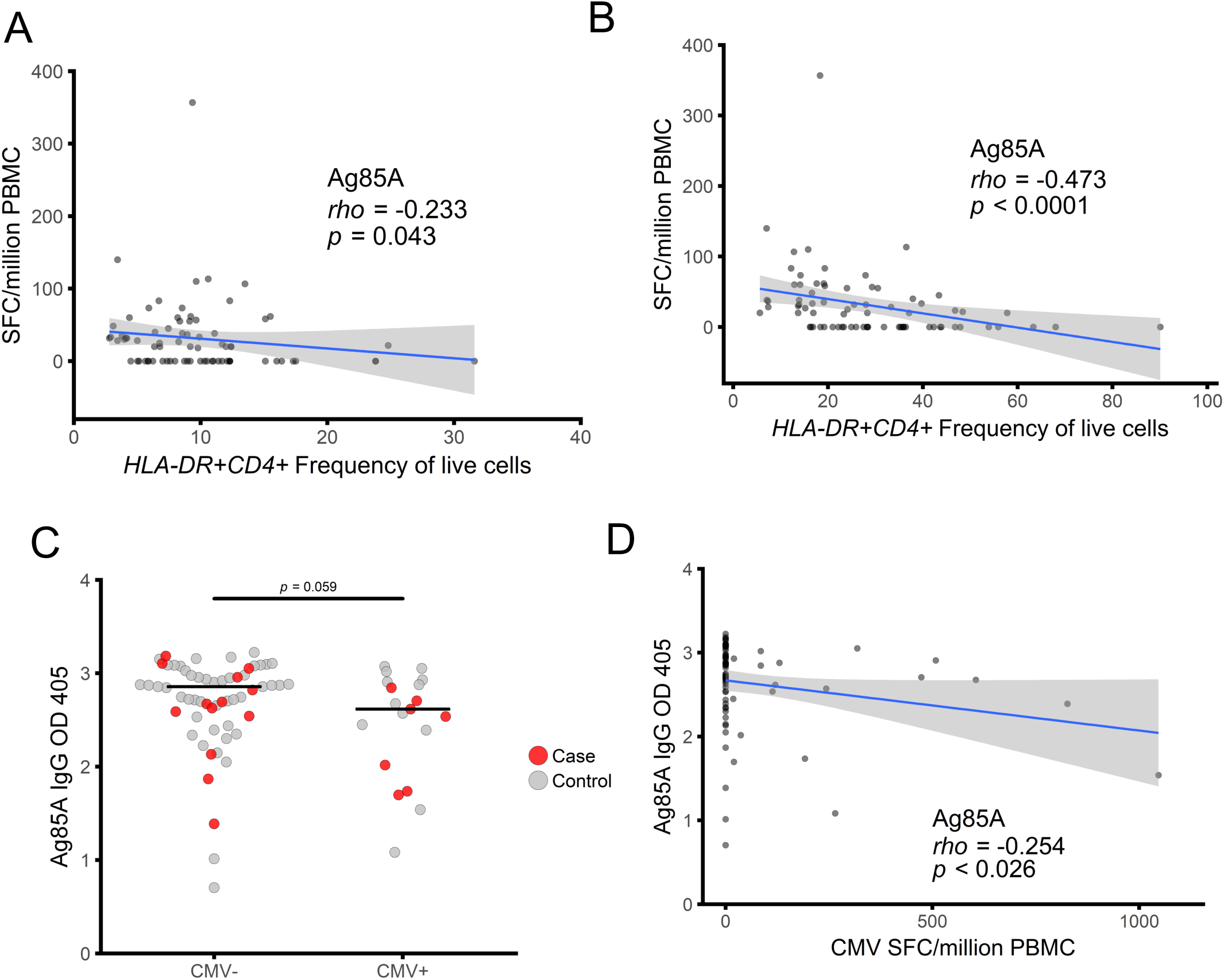
T cell activation is associated with lower mycobacterial antigen specific immune response following immunization with MVA85A. The D28 IFN-*γ* ELISpot response to Ag85A was inversely correlated with both A) activated CD4 T cell and B) activated CD8 T cell frequency C) There was a trend for lower anti-Ag85A IgG in CMV+ compared to CMV- infants following immunization with MVA85A. D) anti-Ag85A IgG was inversely correlated with CMV ELISpot response. Mann Whitney U test and Spearman Spearman’s Rho correlation. Red = cases and grey = controls.

These data show that T-cell activation and CMV infection influence MVA85A boosting of an antigen specific immune response.

## DISCUSSION

We demonstrate, in healthy infants, that CD8+ T-cell activation and prior or sub-clinical infection with CMV (defined by a positive CMV-specific T-cell response) are associated with increased risk of developing TB disease over the next 3 years of life. We also show that CMV+ infants acquire TB disease earlier than CMV- infants. These data complement our previous finding that CD4+ T-cell activation was associated with TB disease risk in infants and adolescents from a South African community with very high TB incidence ^8^. Previous studies have reported that infection with CMV enhances the risk of HIV acquisition and disease progression, through expansion of activated CD8+ T-cells, depletion of naïve T-cells and T-cell senescence ^11-14^ More recently, CMV has been implicated in the aetiology of TB, supported by epidemiological associations between the two diseases ^17-21^. In Gambian infants, CMV infection induced profound CD8+ T-cell differentiation and activation which persisted up to 2 years after infection ^31,32^. Consistent with this effect, we show that infants with a positive T-cell response to CMV peptides have a transcriptional signature associated with CMV specific CD8+ T-cells ^22^ However, among CMV+ infants, T-cell activation markers were not differentially expressed between case and control infants.

In CMV+ infants, transcripts associated with NK cells had lower expression and NK cell frequency was lower in cases when compared to controls. A role for NK cells in protection from TB disease has been demonstrated both in humans and animal models and it is possible than an impaired NK cell response is associated with TB disease risk among CMV+ infants ^33-39^. CMV infection promotes the expansion of NK cells expressing the CD94/NKG2C activating receptor, and these cells are important for control of viral replication ^40^. The CD94 and NKG2C transcripts KLRC1 and KLRC3 had among the greatest decreases in fold-change of expression and the highest predictive power in identifying cases among CMV+ infants. The NKG2C receptor is encoded by the *KLRC2* gene, heterozygous and homozygous deletion of which is present in a significant proportion of individuals across different populations ^41^. *KLRC2* deletion is associated with reduced numbers of mature NK cells and increased susceptibility to HIV infection, certain autoimmune conditions and cancer ^41-43^. We hypothesize that increased susceptibility to TB in CMV+ infants may result from loss of control of CMV replication and/or impairment of NK cell function due to *KLRC2* gene deletions in some individuals ^17^.

Among CMV- infants, who went on to develop TB, we observed upregulation of transcripts associated with T-cell activation, including LAG3 ^23^ which is induced during active TB in a non-human primate model ^44^. We also found multiple transcripts and pathways that are typically altered during viral infection among CMV- infants. This may be due to underestimation of the prevalence of viral infection, as we used only IFN-*γ* responsiveness to EBV and CMV CD8 epitopes as a measure of viral infection. Based on previous studies of viral prevalence in infants in Africa we would expect a CMV prevalence of 24-41% ^36,37^ and EBV prevalence of 35% by 6 months of age ^47,48^. Detection of viral DNA would allow more accurate diagnosis of viral infection, however, this was not possible due to the very limited samples collected from these infants. Moreover, viruses other than those measured in this study (CMV, EBV) may be contributing to risk of TB disease among these infants. Expression of a broad range of both activating and inhibitory KIR receptor transcripts was elevated in CMV- case infants. Exposure to multiple viral infections drives high diversity of KIR expression and lowers the availability of naïve NK cells to respond to future infectious challenge, resulting in susceptibility to HIV ^49,50^. We were able to identify different classifier signatures among CMV+ and CMV- infants and were able to verify our CMV- signature in an independent cohort of infants ^3^. In the independent cohort, infants were younger (2-3 months of age) and few infants were classified as CMV positive. CMV infection and viral replication is low at birth, peaks at 3-6 months of age and declines to plateau at 8-10 months of age ^51^. At 4-6 months of age, infants recruited in to the MVA85A efficacy trial ^1^ were at the peak age of CMV viral replication in infancy.

Our transcriptional evidence of antiviral immune responses in infants who develop TB disease is also consistent with the observations of Zak *et al.* ^2^ who reported increased expression of Type I/II IFN associated transcripts in M.tb-infected adolescents who progressed to TB disease. Increased Type I/II IFN transcripts have also been observed in patients with active TB disease, when compared to *M.tb*-infected or uninfected controls ^52-56^. Recent data from the *M.tb*-infected adolescent study has shown that an increased Type I/II IFN response precedes a shift towards an elevated monocyte to lymphocyte ratio and an increase in T cell activation, which are detected closer to the time of TB disease ^57^. The authors suggest that the initial elevation in Type I/II IFN could be driven by viral infection and that this could then trigger the immune events that lead to TB susceptibility ^57^. Consistent with this we found a correlation of the adolescent CoR score with an EBV response and with elevated CD14+CD16+ inflammatory cells. We also observed significant enrichment of the 16-gene CoR signature identified by Zak *et al.* among our CMV- case infants^2^. However, the CoR signature was not able to accurately classify case and control infants. In our study, the infants were not infected with *M.tb,* had no symptoms of TB disease and no known exposure to TB in the household during enrolment or when the blood samples were obtained. In addition, since incident TB disease was diagnosed months to years after blood sample collection it is unlikely that the elevated type I/II IFN associated transcriptomic signatures we observed are a result of sub-clinical TB disease. Although there was no evidence of BCG vaccine-induced disease in these infants we cannot rule out that persistent, sub-clinical, replication of BCG vaccine could be driving a type I/II IFN response in some infants.

All infants in this study received BCG at birth and the T cell response to BCG peaks at 2-3 months of age ^58^ Viral infections during the development of the BCG-specific immune response may impair the development of protective immunity, as has been suggested by studies in Malawi ^59^ In the Gambia, exposure to HIV *in utero* and being born in the wet season, associated with increased respiratory and diarrheal disease, have been shown to impact the BCG antigen specific T-cell response following vaccination ^31,60^. We observed an inverse correlation between CD4+ and CD8+ T-cell activation and antigen specific T-cell responses following immunization with MVA85A, suggesting that T-cell activation may be associated with decreased vaccine boosting in infants. This confirms previous observations where increased immune activation associated with lower immune responses to MVA85A in both infants and adults ^61,62^ It could also explain why immune responses to MVA85A were lower when administered within a week of vaccination with DTwP-Hib and hepatitis B in the Expanded Programme on Immunization (EPI) schedule ^63^.

Childhood TB is difficult to diagnose and treat and improved strategies are needed to control TB in children ^5^ We found that CMV infection was associated with increased risk of developing TB disease in infants and that distinct immune pathways were associated with TB disease risk in CMV+ and CMV- infants. We suggest that viral infection can increase the risk of progression to TB disease and may compromise the immune response to TB vaccines given in infancy. Strategies which include the use of vaccines or antivirals to reduce chronic viral infection in infancy could enhance TB vaccine efficacy, reduce TB risk and help to reduce the global burden of childhood TB.

## METHODS

### Case-control design

BCG-vaccinated infants who were enrolled in an efficacy trial of the candidate TB vaccine MVA85A were included in this study, ClinicalTrials.gov number NCT00953927 ^1^ (Figure 1). This trial was approved by the University of Cape Town Faculty of Health Sciences Human Research Ethics Committee, Oxford University Tropical Research Ethics Committee, and the Medicines Control Council of South Africa. All infants received BCG within 7 days of birth. HIV and *M.tb* uninfected infants without known TB exposure were randomized at 16-24 weeks of age to receive a single intradermal dose of MVA85A or placebo (Candin™, a candida skin test antigen)^1^.

Transcriptomic analysis was performed using PBMC from infants who were included in our previously described case-control study ^8^. Briefly, infants who met the primary case definition for TB disease were included as cases and for each case, three infants were randomly selected from a pool of controls (Figure 1). Infants were included in the control pool if they did not demonstrate *M.tb* infection as defined by a positive QuantiFERON TB Gold In-tube test (Cellestis, Australia); had not received TB treatment and had not received isozianid preventive therapy during study follow-up. Matching was based on gender, ethnic group, CDC weight-for-age percentile (± 10 points), and time on study (± 9 months).

Infant PBMC samples collected from two time points Day 0 (D0) and Day 28 (D28) were combined for testing of both cellular parameters and transcriptional signatures associated with risk of TB disease. Only infants for whom a sample was available at both D0 and D28 were included in the analysis resulting in combined analysis of a maximum of 98 samples from 49 cases and 258 samples from 129 controls (actual numbers in analysis vary per availability of assay data and are listed in Table 1, Figure 1).

### Cell culture

PBMC were retrieved from liquid nitrogen storage, thawed and rested for 2 hours in media containing DNase to aid the removal of debris from dead and dying cells. After 2 hours cells were counted and immediately transferred to cell culture plates for stimulation for RNA extraction, ELISpot assays or measurement of cell-surface markers assessed by flow cytometry. Viability of thawed PBMC was assessed using flow cytometry with Live/Dead Violet stain (Invitrogen). Phytohemagglutinin (PHA) was included as a positive control for cell viability on ELISpot plates.

### RNA processing

Rested PBMC (1 × 10^6^) were resuspended in single wells of 200μl RPMI supplemented with 10% FBS and L-glutamine. Cells were stimulated for 12 hours at 37°C in single wells containing live BCG SSI from pooled vaccine vials (~2 × 10^5^ CFU/ml). After 12 hours of stimulation cells were pelleted and lysed in RNA lysis buffer (RLT, Qiagen). RNA was extracted using the RNeasy Mini Kit (Qiagen) per the manufacturer’s instructions with the following modification; in the first step an equal volume of 80% ethanol was added to cells lysed in RLT buffer, mixed and total volume transferred to an RNeasy column. RNA was quantified by Nanodrop and stored at −80°C until use. Extracted RNA was amplified and labeled with biotin using the Illumina Total Prep kit (Ambion) per manufacturer’s instructions. Amplified RNA was assessed by nanodrop and Bioanalyzer for quantity and quality prior to hybridization. Hybridization to Illumina HT-12 arrays was performed per manufacturer’s instructions. Arrays were scanned using an Illumina iScan machine and data extracted using Genome Studio software.

### *Ex vivo* IFN-γ ELISpot assay

The *ex vivo* IFN-γ ELISpot assay was performed as previously described using a human IFN-γ ELISpot kit (capture mAb -D1K) (Mabtech) ^8^. Briefly, duplicate wells containing 3 × 10^5^ PBMC were stimulated for 18 hours with antigen, PHA or media alone. Antigens included a single pool of Ag85A peptides (2 μg/ml/peptide) (Peptide Protein Research); BCG (2 × 10^5^ CFU/ml (Statens Serum Institute)); purified protein derivative (PPD) from *M. tuberculosis* (20 μg/ml) (Statens Serum Institute); peptide pools containing known CD8+ T-cell epitopes from EBV (15 peptides), and CMV (5 peptides), (2 μg/ml/peptide, ANASPEC). Results are reported as spot-forming cells (SFC) per million PBMC, calculated by subtracting the mean of the unstimulated wells from the mean of antigen wells and correcting for the number of PBMC. A response was considered positive if the mean number of spots in the antigen well was at least twice the mean of the unstimulated wells and at least 5 spots greater.

### Cell surface flow cytometry

As previously described ^8^ PBMC were washed and stained with 5μl Live/Dead Violet (Invitrogen) followed by surface staining with the following titrated antibodies: 0.5μl CD3-AF700 (clone UCHT1, Ebioscience), 2μl CD4-APC (clone RPA-T4, Biolegend), 2μl CD8-Efluor605 (clone RPA-T8, Ebioscience), 2μl CD14-PE/Cy7 (clone HCD14, Biolegend), 2μl CD16-AF488 (clone 3G8, Biolegend), 1μl CD19-PE/Cy5 (clone HIB19, Biolegend), 2μl CD25-APC/Cy7 (clone BC96, Biolegend), 2μl CD127-NC650 (clone eBioRDR5, Ebioscience) and 15μl HLA-DR-PE (clone L243 Biolegend). Fluorescence minus one (FMO) controls were used to set gates for CD25, CD127 and HLA-DR. Samples were acquired on a BD LSR II flow cytometer. Results are presented as percentages of cells after excluding dead cells and doublets. CD4+ and CD8+ T-cells were identified as CD3+ cells, while CD14+/− and CD16+/− cells were identified as CD3-and CD19-populations. CD25+ CD27-populations were gated on the CD4+ cells. The network representation of cell populations positively correlated among all infants was done using the igraph package in R. To identify closely related clusters (communities) within the network, the ‘cluster_optimal’ function was used implementing an algorithm described in Brandes *et. al.*^*4*^

### Transcriptional analysis

Raw, probe level summary values exported from Illumina GenomeStudio 2011 of Illumina HumanHT 12 V4 microarrays were imported into R using beadarray ^64^. Probes were background corrected using negative control probes followed by quantile normalization using the neqc command ^65^. The analysis was restricted to probes with a detection p-value of <0.01 in at least 10% of the samples and probes matching to the transcript definition of the following databases (in descending importance) with at most 2 mismatches, no insertions and a minimum mapping length of 40 bases: GENCODE version 23, RefSeq (refMrna.fa) and GenBank (mrna.fa) downloaded in August 2015 from http://hgdownload.cse.ucsc.edu/goldenPath/hg38/bigZips/.

A linear model was fitted using limma ^66^ to determine differential expression adjusted for vaccine, day, stimulus, gender, age, ethnicity and batch effects. Array quality weights were incorporated ^67^ to account for between array quality differences. To account for between patient correlations, the duplicateCorrelation command from the limma package was used. Nominal p-values were corrected for multiple hypotheses testing using the Benjamini-Hochberg procedure ^68^. Due to the heterogeneity of the samples, a lenient cut off at an FDR of 20% was chosen to identify genes as significantly differentially expressed. In total 221, 101 and 16 probes mapping to 203, 95 and 16 genes were significantly differentially expressed between Case and Control infants within CMV-, CMV+ and the combined group respectively. For the comparison of EBV+ versus EBV-, CMV+ versus CMV- and CMV strongly positive (ELISpot >100 SFC/million) versus CMV- in total 334, 14 and 103 probes mapping to 296, 14 and 97 genes were significantly differentially expressed respectively. Gene set enrichment analysis was carried out using the cerno test from the tmod package in R. Modules for the enrichment analysis were taken from Li et al.^69^. Datasets used for comparative analysis to CMV infected CD8 T-cells were obtained from Gene Expression Omnibus by downloading GSE12589 and GSE24151.

### Classification

For classification into cases and control samples, we used normalized microarray intensities adjusted for scan date, sample collection time point (Day 0 or Day 28) and stimulation (unstimulated or BCG stimulated). Features for training were selected by using the significant probes at 20% FDR and selecting only the probe with the highest average expression per gene giving the final transcriptional signature size of 95, 203 and 16 genes for CMV+, CMV- and all infants, respectively. Each infant was represented either by sets of two samples (unstimulated and BCG stimulated either at Day 0 or Day 28) or by sets of four samples (unstimulated or BCG stimulated at Day 0 and Day 28). To avoid overfitting, we implemented a modified nested cross validation scheme such that only complete sample sets per infant were assigned to either test or training set at each splitting iteration during the cross-validation process.

Model training was performed using a neural network model as implemented in the nnet package through the caret interface in R ^70^. In the outer loop, samples were split 50 times with replacement into training (about 70%) and test sets (about 30%) to evaluate model performance and feature importance. For model parameter tuning in the inner loop, each training set was split into training and validation sets using a leave-one-infant-out cross validation scheme and the AUROC was recorded as performance metric over a grid of 4 size and 4 decay parameter combinations. All accuracies stated in the manuscript are provided as balanced accuracies^71^ to account for the imbalance of the case and control infants within our cohort.

To validate our classification results, an independent cohort of 10-week-old South African infants vaccinated with BCG at birth was obtained as raw expression data from Gene Expression Omnibus by downloading GSE20716. Study batch was removed with the ComBat command in R using parametric adjustment, risk of TB as null model and the validation cohort as reference in order to avoid any bias on the validation cohort, For prediction of risk of TB, 112, 50 and 5 probes overlapped between Illumina HumanHT 12 V4 and Illumina HumanRef-8 V2 for within CMV-, CMV+ and the combined group respectively. CMV status prediction was performed using 55 overlapping probes which were differentially expressed between CMV negative and CMV strongly positive (ELISpot >100 SFC/million) infants at an FDR of 20%. Only infants with at least two positive samples were labelled as suspected CMV+.

## Data availability

Raw and normalized expression data have been deposited at Gene Expression Omnibus under the accession number GSE98550

## Acknowledgements

We thank study participants and their families, the community of Cape Winelands East district, and South African Tuberculosis Vaccine Initiative (SATVI) personnel. This work was funded by Aeras and The Wellcome Trust with support from the European Commission within the 7th framework program (FP7) NEWTBVAC (Grant No. HEALTH-F3-2009-241745) and by the European Commission within Horizon2020 TBVAC2020 (Grant No. H2020 PHC-643381). HM is a Wellcome Trust Senior Clinical Research Fellow.

---

**Additional Figure 1.**
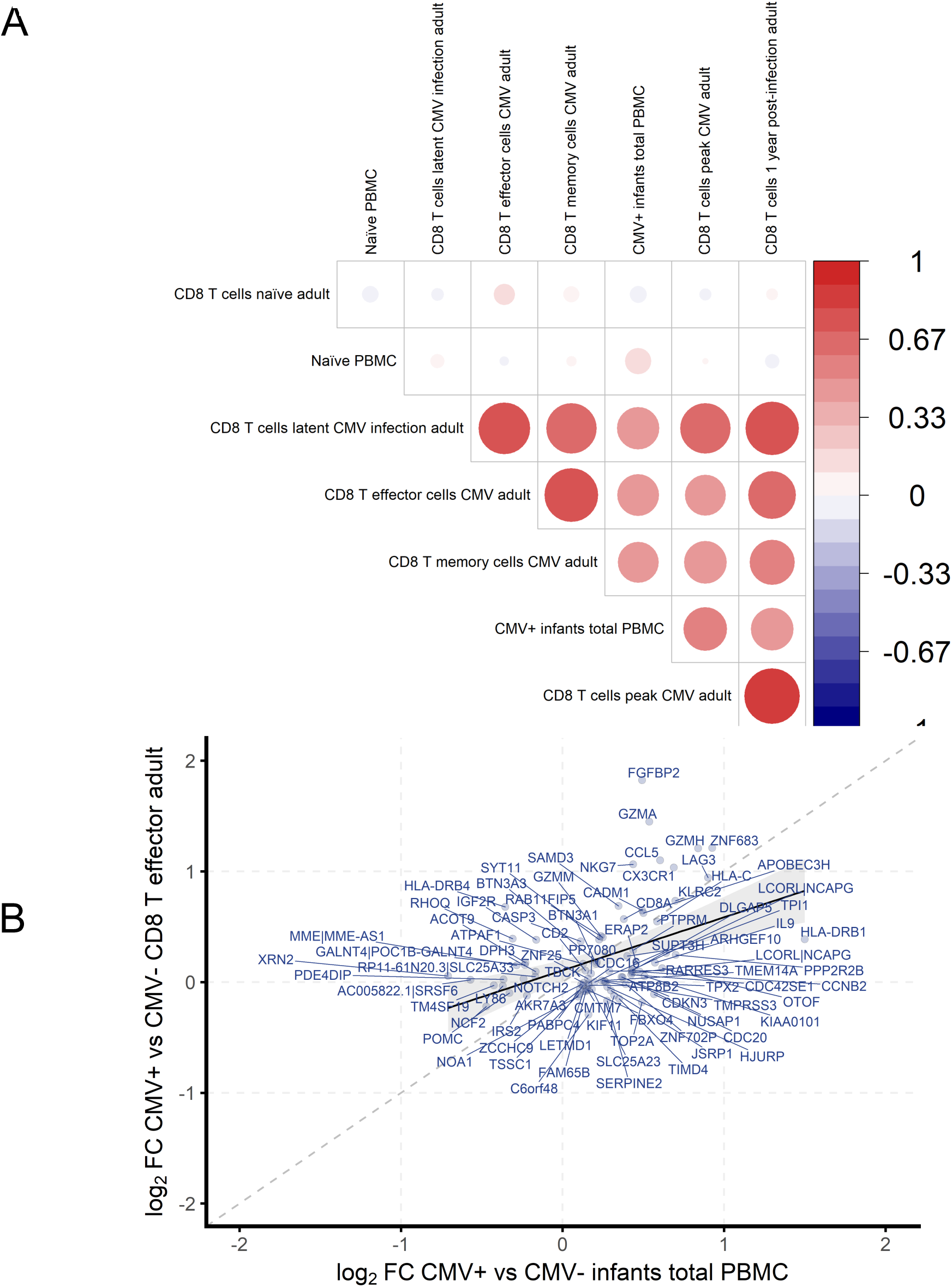
Prominence of CD8 T cell specific transcripts from total PBMC of CMV ELISpot positive infants. A) Spearman’s rho from correlations of gene expression from total PBmC of infants with a strong CMV ELISpot response >100 SFC/million with CD8+ T cells isolated from adults who were naïve or infected with CMV (transcripts with <20% FDR). B) Scatter plot of 84 overlapping transcripts between PBMC of infants with a strong CMV ELISpot response >100 SFC/million and transcripts from CMV virus specific effector CD8 T cells from CmV infected adults (GSE24151).

**Additional Figure 2.**
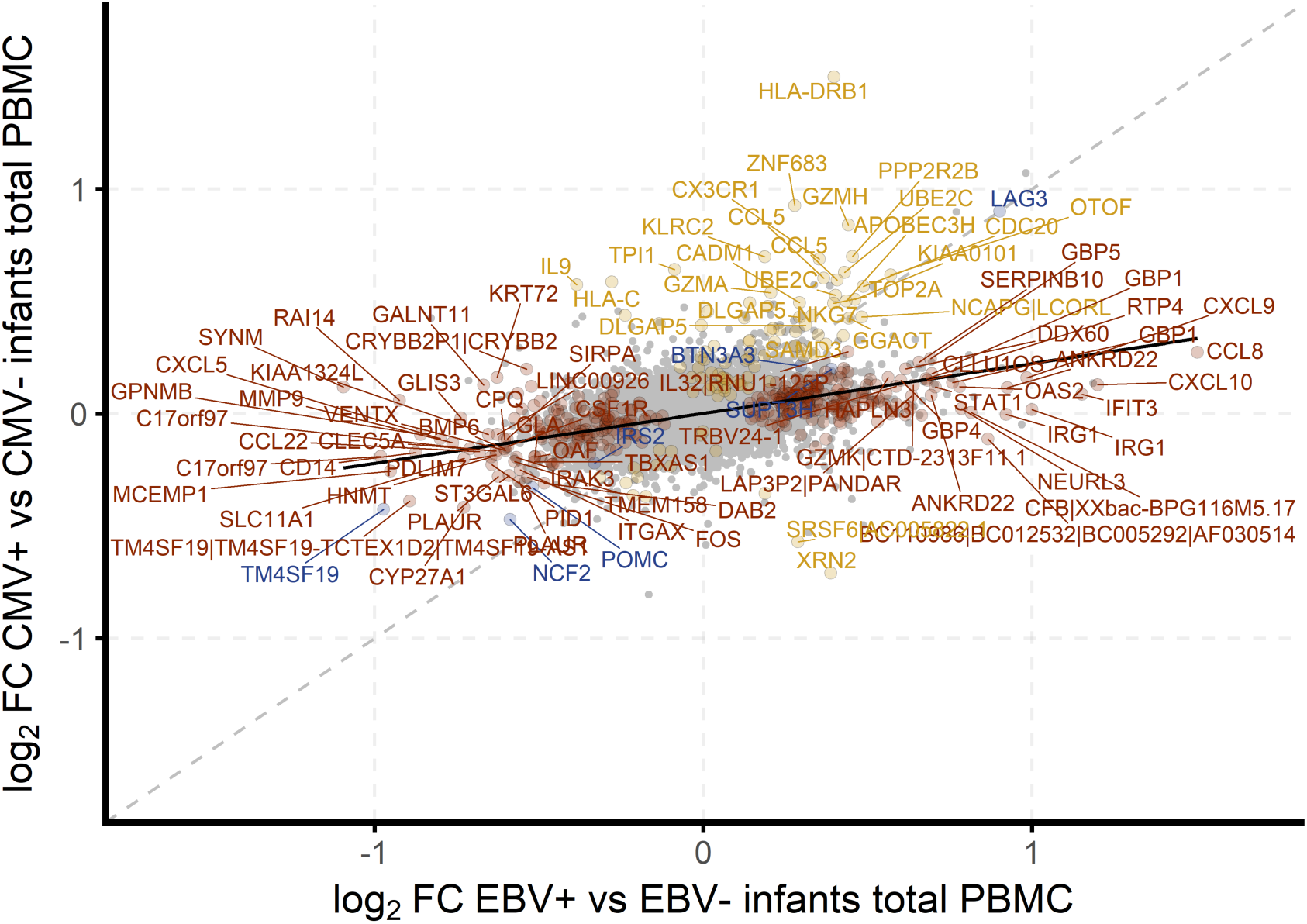
Little overlap in expression in infants who are either EBV or CMV ELISpot positive. Scatterplot of fold changes in response to EBV infection (x-axis) plotted against changes in response to CMV infection (y axis). The almost horizontal linear regression line indicates stronger changes in response to EBV. At an FDR of 20%, seven genes are significantly differentially expressed in both comparisons.

**Additional Figure 3.**
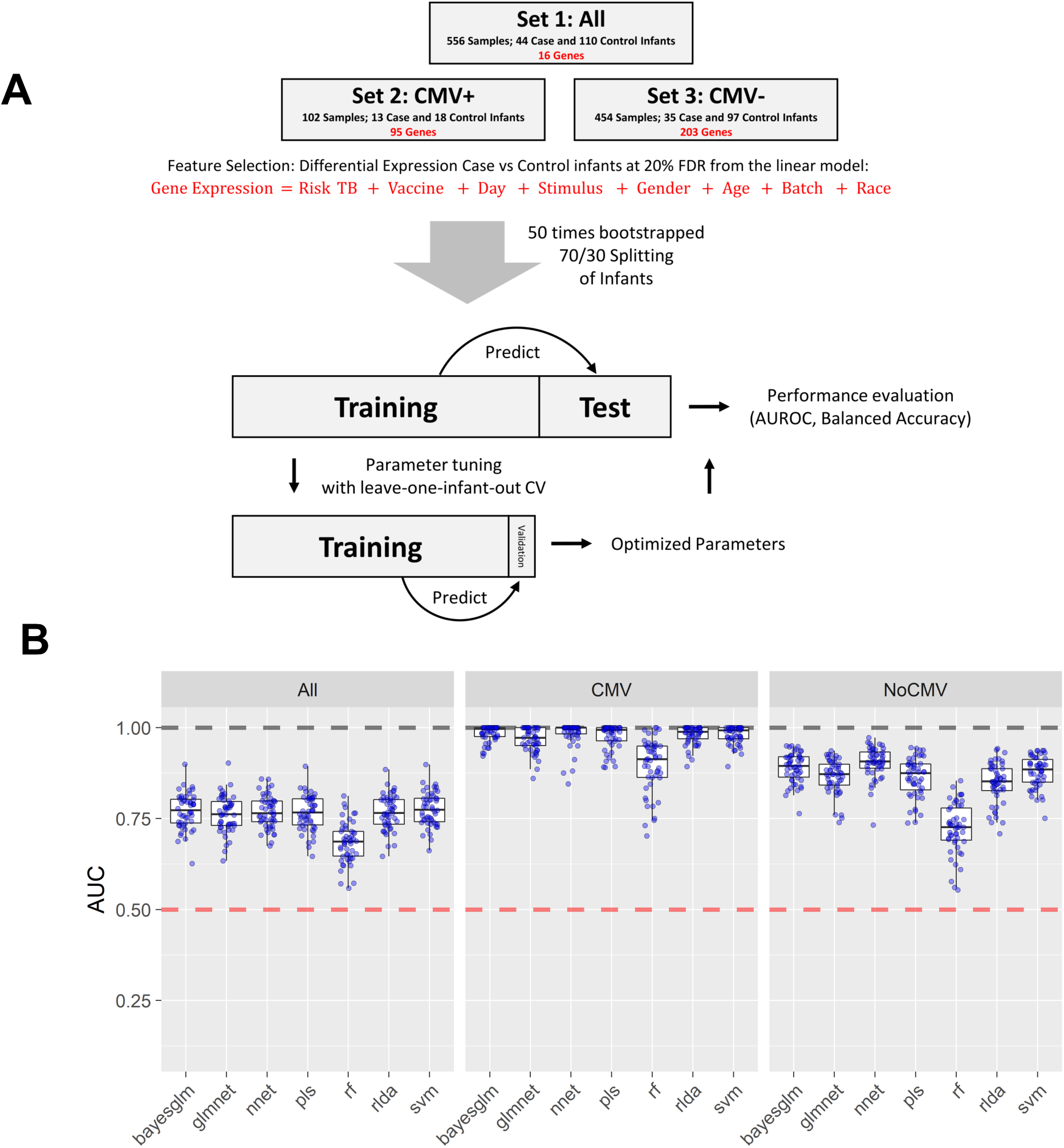
Classification strategy and model comparison. A) Overview of the classification strategy for TB risk prediction. An artificial neural network model was trained using approximately 70% of the data and TB risk was predicted on the withheld 30% of the data. Only significantly differentially expressed genes were used as features for classification for each set of infants and model parameters were tuned using leave one infant out cross validation. The process was repeated fifty times with random splits into training and test set (bootstrapping) and AUROC and balanced accuracy were recorded for each round. B) Classification performance comparison of seven models applied to the datasets. Full names of models and parameter tuning as follows: bayesglm: Bayesian Generalized Linear Model, no tuning parameters; glmnet: a grid of 4 lambda values as determined by a call to glmnet and a fixed alpha at 0.1 to enforce moderate feature selection; nnet: Neural Networks, grid of two unit size (1 and 3) and 3 decay (0.1, 1 and 2) parameters.; pls: Partial Least Squares, grid of 10 components; rf: random forrest, grid of 20 mtry values; rlda: Regularized Linear Discriminant Analysis with Schafer-Strimmer estimator, no tuning parameters; svm: Support Vector Machine with linear kernel, grid of 10 C values

**Additional Figure 4.**
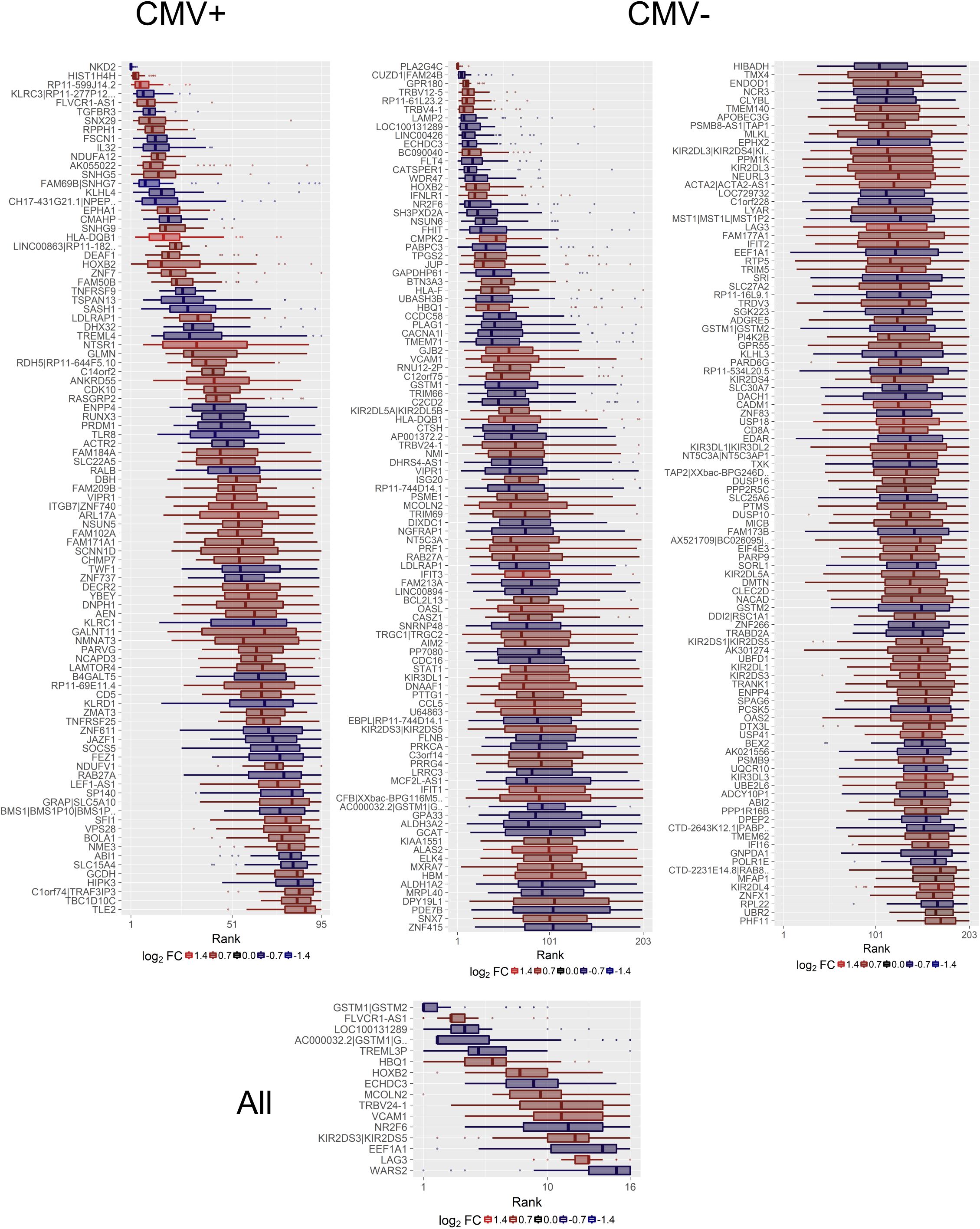
Boxplots showing the relative importance of features used to train the classifier for each data set. For each repeat, features were ranked by variable importance and the relative rank was recorded. The lower the average number, the more often a feature has been assigned a high importance by the classifier during the fifty repeated predictions. The color indicates the Case vs Control infants log2 fold change estimate based on the differential expression analysis

**Additional Figure 5.**
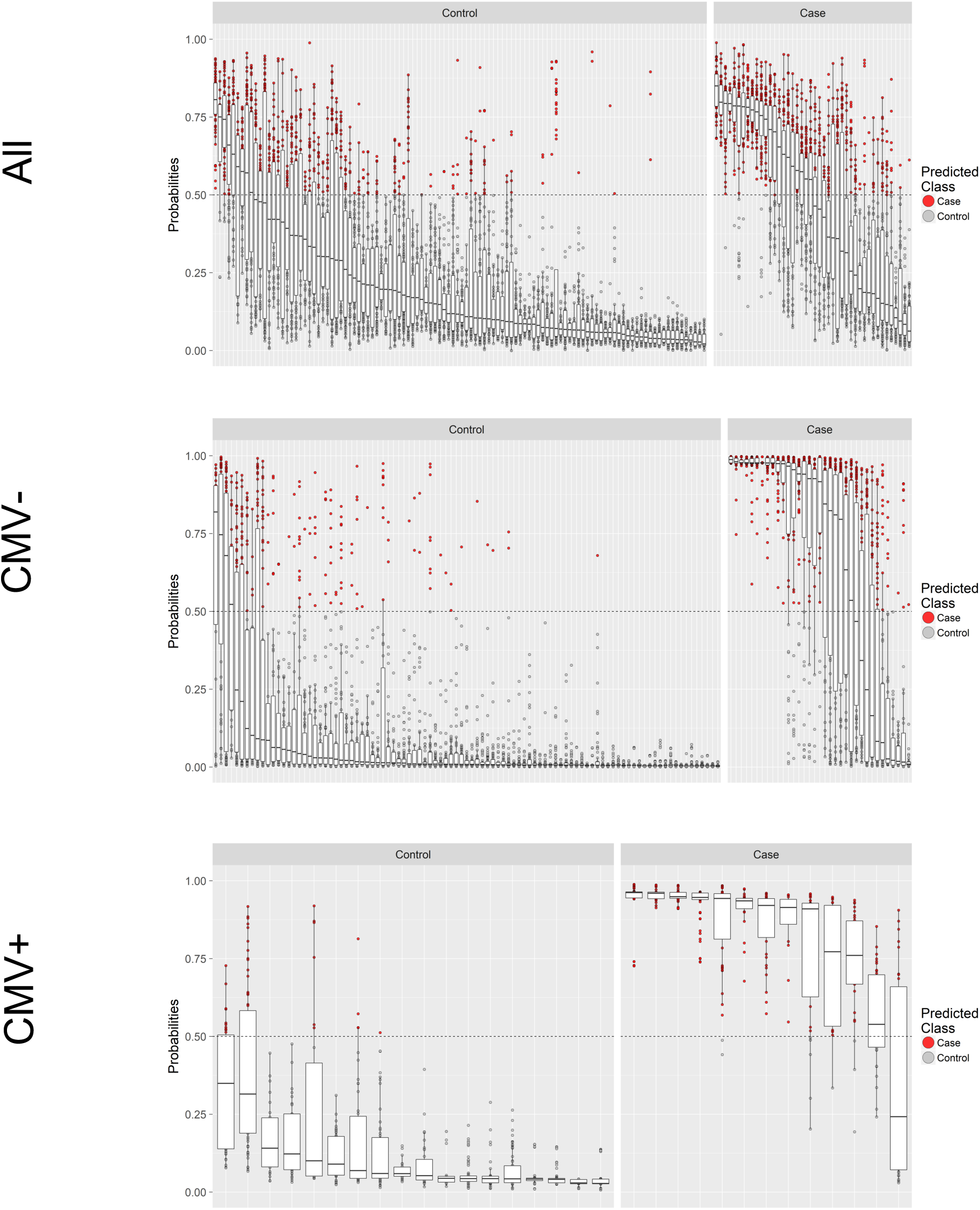
Average prediction accuracies for risk of TB are very high in infants stratified by their CMV status. Box plots of probabilities assigned to each sample by the trained neural network. Each box plot represents one infant and contains stimulated as well as unstimulated samples and the results of fifty bootstrapped repeats of the prediction. Left and right panel divide the infants into cases and controls and point color indicates the classification decision by the model.

**Additional Figure 6.**
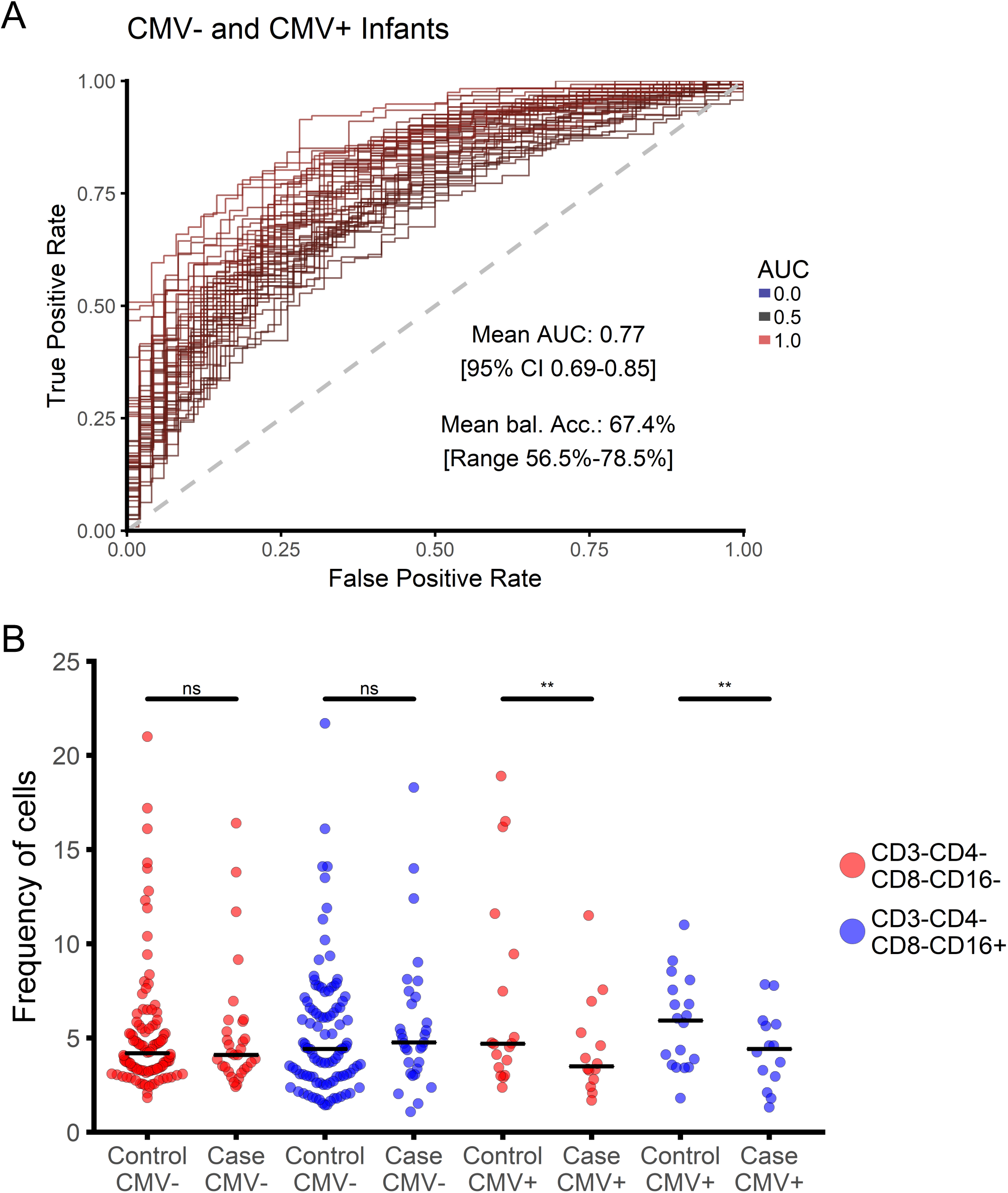
Classification into cases and controls using all samples. A) Classification performance of the artificial neural network using CMV+ and CMV- infants. B) CD16- and CD16+ (putative) NK cell frequencies among case and control infants with and without a CMV response

**Additional Figure 7.**
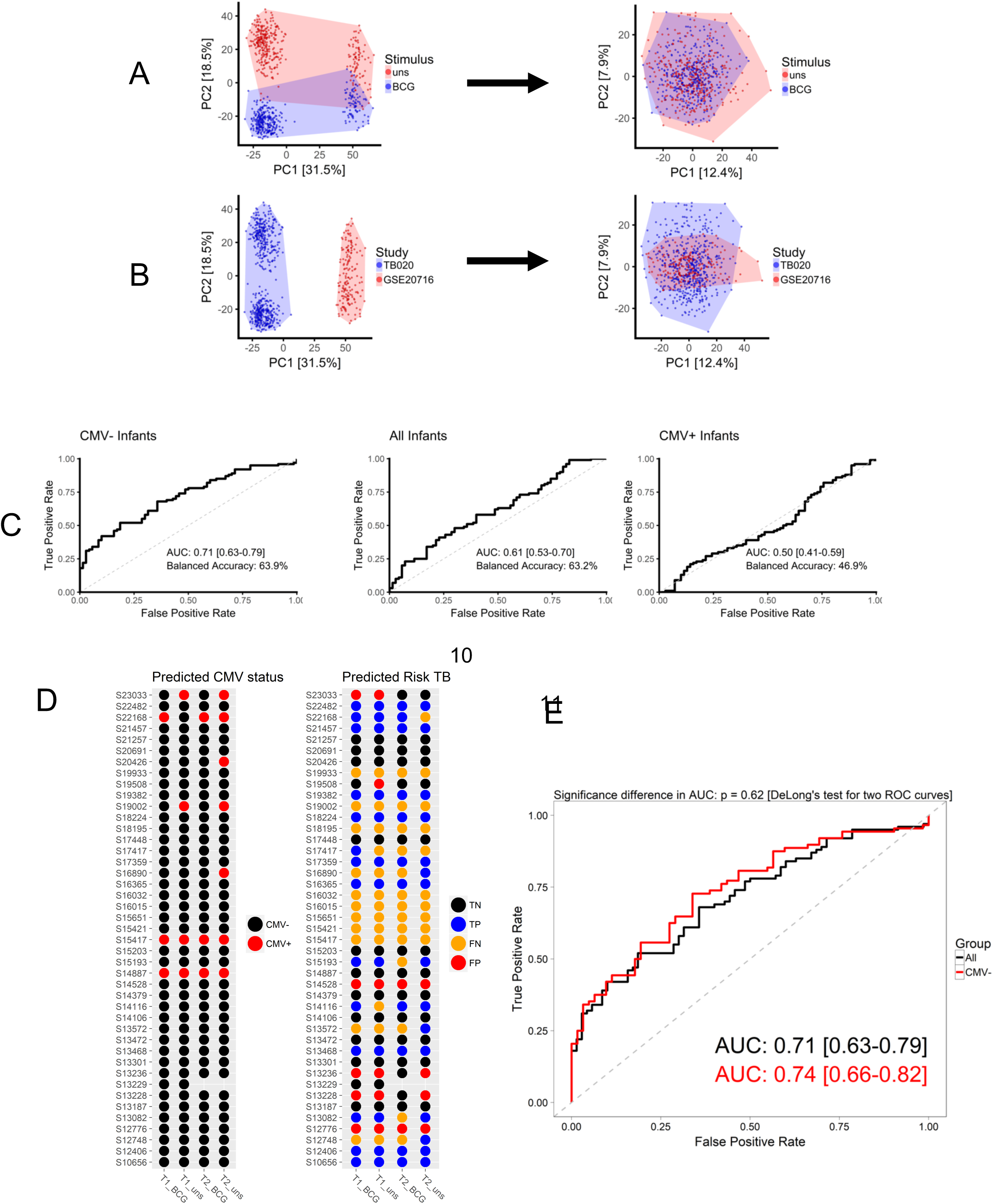
Validation of TB predictive biomarkers from CMV- infants in an independent cohort of BCG vaccinated infants. Raw data and detection p values from an independent cohort of 10 week old South African infants vaccinated with BCG at birth (GEO gene set GSE20716^3^) were used as a validation cohort for the classifier signatures from CMV+ and CMV- infants. 20589 Probes from 182 samples were mapped from Illumina HumanRef-seq 8 arrays to Gencode v.23 and 8 outlier samples were excluded A) The effects of time point (4 or 12 hours) and BCG Stimulation (stimulated or unstimulated) were removed and 9195 probes identified as overlapping between the two studies. B) We were only partially able to remove differences in gene expression due to study of origin. The differences observed in PC2 are driven by age, which was non-overlapping in the two studies, 2-3 months in GSE20716 and 4-6 months in the MVA85A efficacy trial.C) Using 55 genes differentially expressed between infants with very high or low CmV titres at an FDR of 5%, 5 infants were identified to have two or more CMV positive samples. D) The charts represents one dot per sample and red colour indicates samples classified as CMV+. The TB Risk dot plot shows samples with correctly predicted Case (TP) and Control (TN) status as well as samples with wrong Case (FP) and Control (FN) labels. E). Excluding all five suspected CMV+ infants we were able to improve risk of TB prediction accuracyon GSE20716^3^slightly, with balanced accuracy of 65.6%.

**Additional Figure 8.**
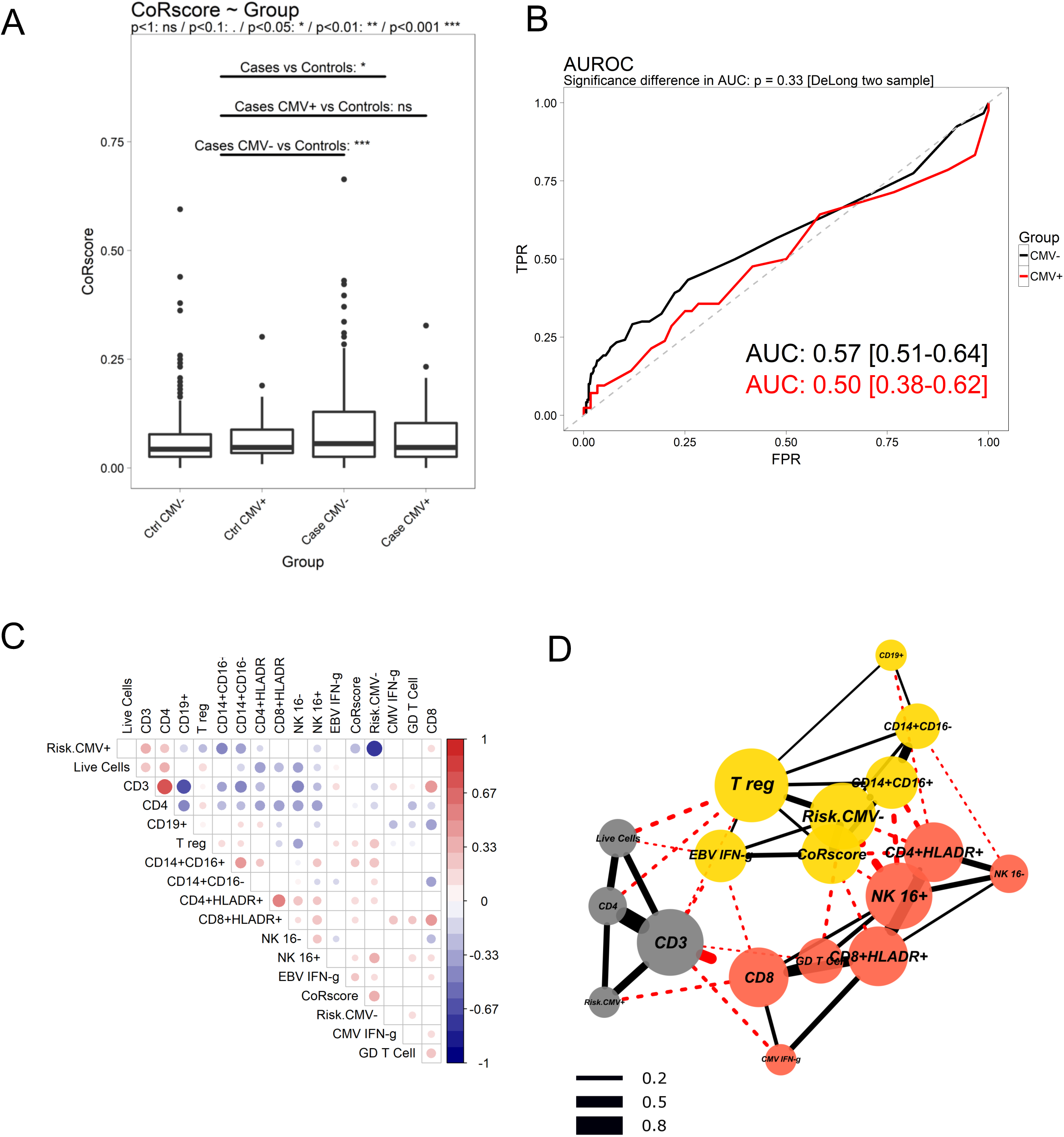
Enrichment of TB predictive Correlate of TB Risk score (CoR) from adolescents among CMV- case infants when compared to controls, but no accuracy for prediction of TB in infants. Using a locked-down Illumina model published by Zak et al^2^ for prediction of TB progression, a blinded CoR score was derived for each infant. A) After unblinding, the CoR score was found to be significantly higher in case infants when compared to controls and the greatest difference in CoR score was between CMV- case and control infants B) However, despite enrichment in case infants, the CoR score was unable to accurately classify either CMV+ or CMV- infants as cases or controls. C) The CoR score correlates with the frequency of inflammatory monocytes and the CMV- infant classifier signature. D) Network of positively correlating cell populations (spearman rho p-value <0.05) showing a cluster containing the adolescent CoR score with monocytes, EbV ELISpot and the infant CMV- classifier score. Node colour indicates community membership and red and black edges are drawn between and within communities respectively (see methods). Edge width indicates the correlation coefficient.

